# Hda1C restricts the transcription initiation frequency to limit divergent non-coding RNA transcription

**DOI:** 10.1101/2021.04.06.438606

**Authors:** Uthra Gowthaman, Maxim Ivanov, Isabel Schwarz, Heta P. Patel, Niels A. Müller, Desiré García-Pichardo, Tineke L. Lenstra, Sebastian Marquardt

## Abstract

Nucleosome-depleted regions (NDRs) at gene promoters support initiation of RNA Polymerase II transcription. Interestingly, transcription often initiates in both directions, resulting in an mRNA, and a divergent non-coding (DNC) transcript with an unclear purpose. Here, we characterized the genetic architecture and molecular mechanism of DNC transcription in budding yeast. We identified the Hda1 histone deacetylase complex (Hda1C) as a repressor of DNC in high-throughput reverse genetic screens based on quantitative single-cell fluorescence measurements. Nascent transcription profiling showed a genome-wide role of Hda1C in DNC repression. Live-cell imaging of transcription revealed that Hda1C reduced the frequency of DNC transcription. Hda1C contributed to decreased acetylation of histone H3 in DNC regions, supporting DNC repression by histone deacetylation. Our data support the interpretation that DNC results as a consequence of the NDR-based architecture of eukaryotic promoters, but that it is governed by locus-specific repression to maintain genome fidelity.

## INTRODUCTION

RNA polymerase II (RNAPII) transcribes DNA of eukaryotic genomes into RNA molecules (Osman & Cramer, 2020). Widespread transcriptional activity of RNAPII in non-protein-coding regions of genome results in non-protein-coding RNAs (David *et al*., 2006; Kapranov *et al*., 2007; Core *et al*., 2008). The transcriptional noise hypothesis posits that non-coding transcription reflects inaccuracies of cellular machinery that carry little functional significance (Struhl, 2007; Ponjavic *et al*., 2007). However, co-transcriptional events resulting from the act of non-coding transcription impact cellular functions ranging from gene regulation to DNA replication (Gowthaman *et al*., 2020). Although the purpose of genome-wide non-coding transcription remains elusive, a growing number of cellular functions carried out by non-coding RNAs (ncRNAs) (Quinn & Chang, 2016; Gil & Ulitsky, 2020) and the dedicated regulation of non-coding transcription (Jensen *et al*., 2013), indicates that interpreting non-coding transcription through the transcriptional noise hypothesis may be an oversimplification.

Divergent non-coding (DNC) transcription from eukaryotic gene promoter regions (Seila *et al*., 2008; Neil *et al*., 2009; Xu *et al*., 2009; Ntini *et al*., 2013) exemplifies the tight regulation of non-coding transcription since transcription initiates in both directions at nucleosome-depleted regions (NDRs) that characterize eukaryotic promoters. DNC transcription is characteristic of a wide range of eukaryotic organisms (Wang *et al*., 2009; Neil *et al*., 2009; Sigova *et al*., 2013; Core *et al*., 2014). Interestingly, the strength of DNC transcription is variable and restricted more tightly in plants, compared to yeast and mammals (Kindgren *et al*., 2020). Cellular pathways that selectively activate or repress DNC transcription may operate with variable intensities and thereby account for the differences in DNC expression across organisms.

The budding yeast has a high-density genome and relatively high levels of DNC transcription, which results in a high likelihood of DNC transcription invading neighboring genes to affect gene expression and fitness (Xu *et al*., 2011; Bumgarner *et al*., 2012; Alcid & Tsukiyama, 2014; Du Mee *et al*., 2018; Moretto *et al*., 2018). However, since functional DNC transcription often results in relatively short-lived RNA species elucidating the cellular functions of DNC remains a challenge. DNC transcripts are targeted for co-transcriptional RNA degradation by the nuclear exosome pathway (Xu *et al*., 2009) and this mechanism of DNC repression is conserved in mammals (Almada *et al*., 2013; Ntini *et al*., 2013). Experimental detection of DNC transcription is thus facilitated by analyses of nascent RNAPII transcription (Churchman & Weissman, 2011) and RNA profiling of RNA degradation pathway mutants (Wyers *et al*., 2005; Neil *et al*., 2009; Xu *et al*., 2009; Van Dijk *et al*., 2011; Schulz *et al*., 2013). In addition to selective RNA degradation, the DNC transcript and the corresponding mRNA are differentially regulated at the level of transcriptional initiation by sequence-specific transcription factors (Neil *et al*., 2009; Challal *et al*., 2018; Wu *et al*., 2018) that lead to the formation of distinct pre-initiation complexes that drive transcription in each direction of the NDR boundaries (Rhee & Pugh, 2012). Finally, chromatin-based mechanisms provide a third pathway for regulation (Whitehouse *et al*., 2007; Tan-Wong *et al*., 2012; Marquardt *et al*., 2014; Rege *et al*., 2015). Among the chromatin factors influencing transcription, histone acetylation, which is governed by histone acetyltransferases (HATs) and histone deacetylases (HDACs) (Kurdistani & Grunstein, 2003; Park & Kim, 2020), is of particular interest. In budding yeast, the Hda1 complex (Hda1C) removes acetylation on residues in histones H2A, H2B and H3 (Carmen *et al*., 1996; Wu *et al*., 2001) and Hda1C-linked histone acetylation facilitates dynamic transcriptional responses (Nicolas *et al*., 2018; Chen *et al*., 2019). Specifically, acetylation of histone 3 lysine 56 (H3K56ac) mediates histone exchange of the −1 nucleosome, and disruption of the H3K56ac homeostasis through mutations in HDACs or the Chromatin Assembly Factor I (CAF-I) histone chaperone complex amplifies DNC transcription genome-wide (Marquardt *et al*., 2014; Rege *et al*., 2015). In summary, DNC transcription is highly regulated in budding yeast, yet the full spectrum of regulatory activities and their interconnections remains elusive.

Transcription of a gene occurs by selective initiation of a single RNAPII (constitutive expression) or in bursts, through high activity of several RNAPIIs (Zenklusen *et al.,* 2008). The different mechanisms of initiation account for cell-to-cell variability that result in heterogeneous transcriptomes even in homogeneous growth conditions (Swain *et al.,* 2002). The dynamics of DNC transcription remain unexplored. Single-molecule imaging offers the possibility to record transcription events and estimate the parameters of transcription dynamics. Single-molecule fluorescence *in situ* hybridization (smFISH) allows visualization of transcripts in fixed cells, and the MS2/PP7 dual RNA-aptamer-based reporter system allows the observation of single transcription events in living cells (Hocine *et al*., 2013; Lenstra *et al*., 2015). Recording transcription initiation by live-cell imaging reveals the frequency and duration of transcription initiation, whereby the frequency determines the rate of initiation and the duration indicates RNAPII transition or elongation (Larson *et al*., 2011; Lenstra *et al*., 2015). Ultimately, transcription frequency and duration depend on the active state of promoters, binding of sequence-specific transcription factors and local epigenetic states may modulate the transcription parameters (Donovan *et al.,* 2019; Rodriguez & Larson, 2020). Histone acetylation modulates the burst frequency of mammalian genes (Nicolas *et al*., 2018; Chen *et al*., 2019). However, the interplay between chromatin-based promoter regulation, transcription initiation kinetics, and DNC remains largely unclear.

In this study, we identify Hda1C as a key factor in the selective repression of DNC transcription. We utilize high-throughput genetic screens based on quantitative single-cell biology methods to characterize the effects of all non-essential genes on DNC. Live-cell single-molecule imaging suggests that DNCs are transcribed constitutively rather than in bursts. Hda1C repression of DNC transcription correlates with reduced DNC initiation frequency and deacetylation of H3. Overall, Hda1C limits DNC transcription genome-wide, presumably by restricting the frequency of transcription initiation in the non-coding direction of promoter NDRs.

## RESULTS

### Genetic screens identify Hda1C and SAGA as regulators of DNC transcription

To identify DNC regulation independent of previously identified mechanisms, we focused on *GCG1*/*SUT098* and *ORC2*/*SUT014* DNC loci. These loci are characterized by high levels of DNC transcription and scored as unaffected by CAF-I mutations that increase DNC at any many loci in the genome (Marquardt *et al*., 2014) (Figure S1 A-C). Native Elongating Transcript sequencing (NET-seq) data reveals higher nascent transcription levels in the non-coding direction than the protein-coding gene at these loci in wild type (WT) (Figure 1A) (Churchman & Weissman, 2011; Marquardt *et al*., 2014; Harlen *et al*., 2016; Fischl *et al*., 2017). We created fluorescent protein reporters (FPR) where the coding and DNC sequences of the *GCG1* and *ORC2* promoter region are replaced with mCherry and Yellow Fluorescent Protein (YFP) sequences to estimate transcriptional activity in each direction (Figure 1B). The resulting strains with the FPR inserted in the non-essential *PPT1/SUT129* locus are compatible with high-throughput reverse Synthetic Genetic Array (SGA) technology (Baryshnikova *et al*., 2010; Marquardt *et al*., 2014). We crossed FPR strains with the library of non-essential gene deletion mutants (Winzeler *et al*., 1999) to perform SGA. The resulting haploid strains harbor the FPR and a specific gene deletion arrayed in microtiter plates. We quantified mCherry and YFP for up to 50,000 single cells for each non-essential gene deletion for robust phenotyping by high-throughput flow cytometry (Figure 1C). Later, we processed the median value for mCherry and YFP fluorescence to plot and compare the effects of individual non-essential gene deletions. We performed two key analyses to identify potential DNC regulators. First, we calculated the directionality score for every mutant in each of the two genetic screens (Table S1). The directionality score captures the distance of a data point from the plate regression line (Figure S1D). We categorized the mutants above the regression line with a positive directionality score as repressors of DNC transcription since we expect increased DNC levels in the mutants. Conversely, we categorized those below the regression line with negative scores as activators of DNC transcription (Figure S1D). Second, we focused our analysis on hits shared between the two screens. Here, we compared the overlap of mutants with a statistically significant directionality score in the individual screens (Figure S1E, Table S2). This analysis identified the two major subunits of Hda1C, *hda1*Δ and *hda3*Δ above the regression line in both the screens (Figure 1D, E). In addition, *hda2*Δ scored as a hit for the *ORC2*pr screen, but not for the *GCG1*pr screen, highlighting all three Hda1C subunits as hits. Mutants of other HDACs and factors linked to protein urmylation scored as repressors as well (Figure S1F). Several mutations in the SAGA subunits such as *ada2*Δ, *ngg1*Δ, *sgf73*Δ and *ubp8*Δ clustered below the regression line, suggesting roles in DNC activation (Figure 1D, E). The analysis of the screen data also identified SWI/SNF subunits and ISW2 chromatin remodelers as activators (Figure S1G). Since we identified most non-essential subunits as common hits in two independent genetic screens, we considered the Hda1C and the SAGA histone acetylation complex as promising hits and potential regulators of DNC transcription.

**Figure 1.**
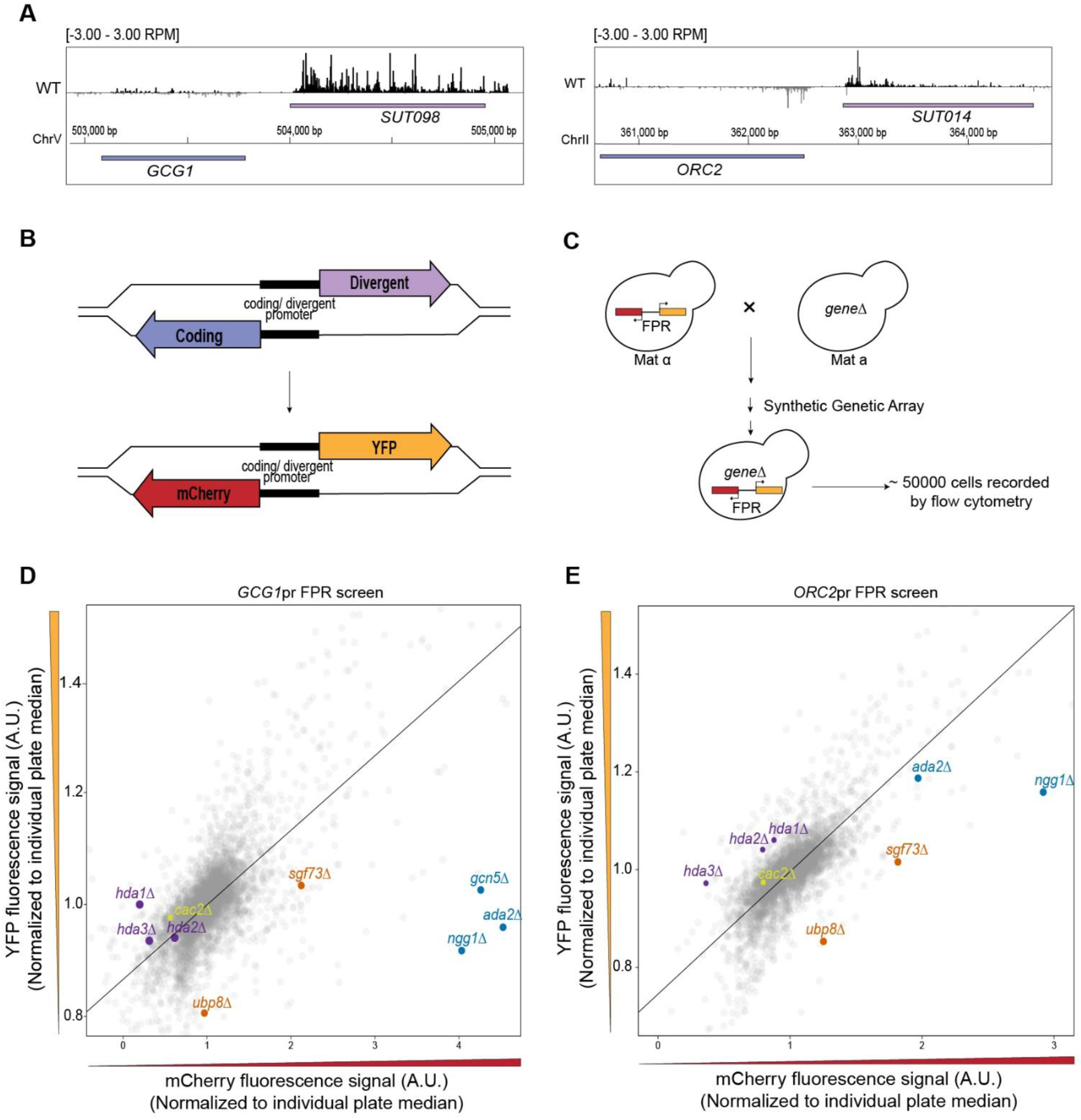
Identification of factors regulating divergent non-coding (DNC) transcription by a synthetic genetic array (SGA) and flow cytometry analysis. (A) NET-seq data of wild type (WT) yeast at the *GCG1/SUT098* and *ORC2/SUT014* loci. NET-seq reads in black and grey represent the Watson (+) and Crick (−) strands, respectively. The NET-seq track combines remapped data from previous publications (Churchman and Weissman., 2011, Marquardt *et al*., 2014, Harlen *et al*., 2016, Fischl *et al*., 2017, see Material and Methods for more details). (B) Schematic representation of a shared promoter region initiating a coding (blue) and a DNC (purple) transcript in the opposite orientation. Fluorescent protein reporter (FPR) construction replaces the endogenous coding and DNC genomic regions with sequences encoding mCherry and YFP, respectively. (C) SGA and flow cytometry analysis in budding yeast. The array involves the crossing of MATα query strain harboring the FPR construct with the MATa non-essential gene deletion mutant library. The systematic selection as described in Baryshnikova *et al*., 2010 results in haploids containing the gene deletion and FPR. Quantification of the resultant haploids by high-throughput flow cytometry records fluorescent signals in up to 50,000 cells for each gene deletion mutant. (D, E) Scatter plot visualization of YFP vs mCherry fluorescence signals in the *GCG1* and *ORC2* promoter FPR screens. Each data point represents the median signal of a deletion mutant normalized to individual plate median. Highlighted data points represent the mutants of Hda1C (purple), SAGA acetylation module (blue), SAGA deubiquitination module (orange), and *cac2*Δ (yellow). The regression line is marked in black. The mutants favoring DNC transcription are found above the regression line. The mutants favoring coding transcription are below the regression line.

To evaluate shared hits between both screens systematically, we asked whether the combined analysis of the screen data identifies other protein complexes. We plotted the directionality scores of each mutant calculated from the two reporter screens to visualize the clustering of mutants with similar attributes. A two-dimensional scatter plot of the directionality score distributes the data points representing gene deletions in four quadrants. We expected the mutants with strong positive or negative directionality scores in both the screens to cluster in quadrant I and III, respectively (Figure 2A). Mutants with high reproducibility between the screens appear closer to the diagonal and those with high directionality score away from the origin. In quadrant I, we observed the Hda1C mutants as strong hits (Figure 2B). We also discovered majority of non-essential subunits of protein urmylation, Rpd3 histone deacetylation, CAF-I, and histones in quadrant I, highlighting their role in DNC repression. In quadrant III, we identify the mutants of the SWI/SNF and ISW2 chromatin remodelers alongside the SAGA complex (Figure 2B). In summary, our high-throughput reverse genetic screens were able to resolve protein complexes likely contributing to the regulation of DNC transcription.

**Figure 2.**
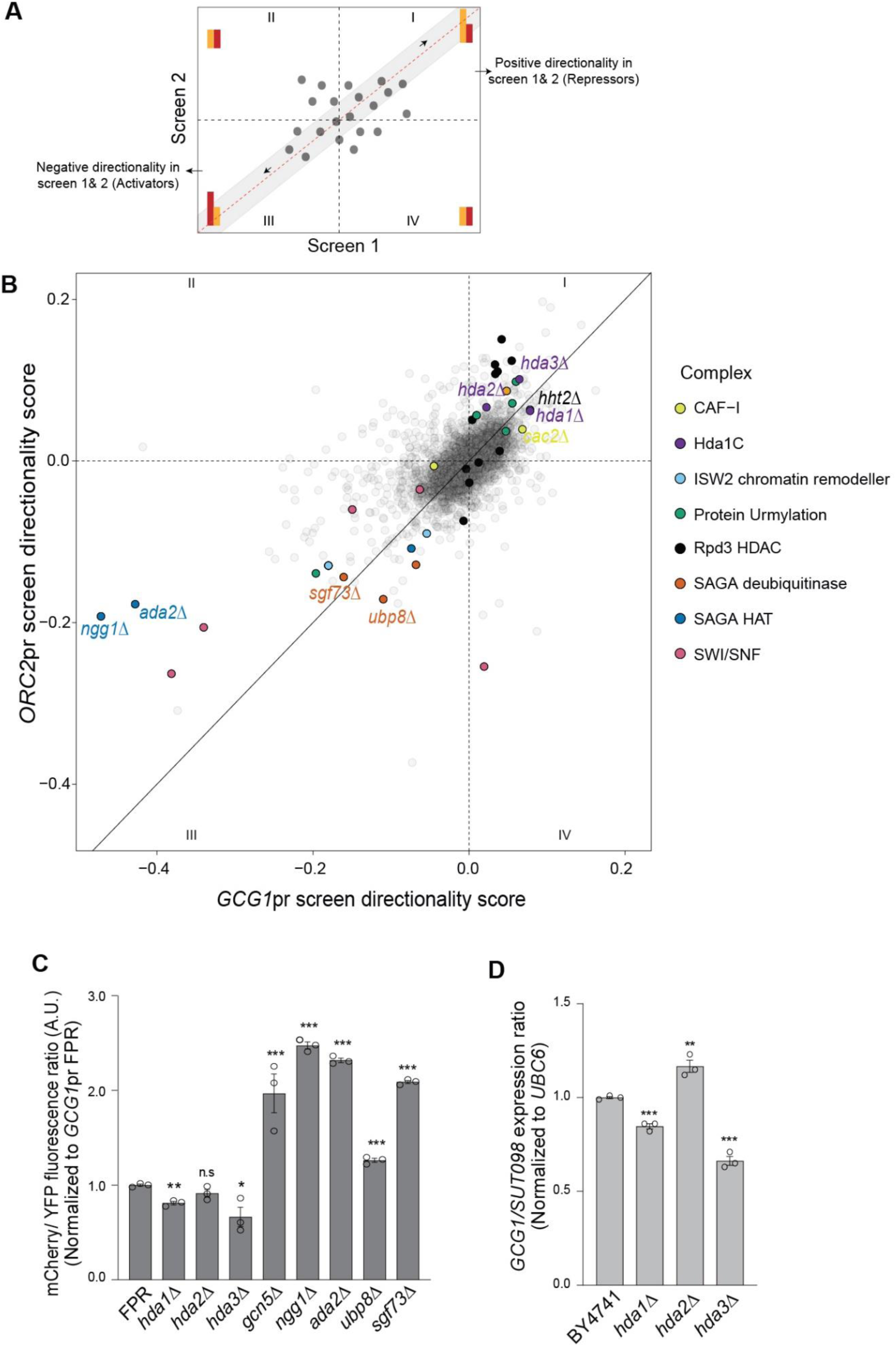
The reverse-genetic screening approach identifies several novel protein complexes with a potential role in the regulation of divergent non-coding (DNC) transcription. (A) Illustration of the expected distribution of data points combined from independent genetic reporter screens. Each data point represents the directionality score of a deletion mutant. Mutants altering directionality positively by increasing divergent YFP levels are expected to be found in quadrant I. Mutants decreasing YFP with negative directionality scores are found in quadrant III. Data points closer to the diagonal (grey region) have high reproducibility between the screens. (B) Distribution of mutants by directionality scores from the *GCG1*pr and *ORC2*pr genetic screens. Quadrant I and III reveal the non-essential mutant subunits of several protein complexes (highlighted in colors) altering the directionality. The top candidate repressors and activators of DNC are labeled. (C) Flow cytometry quantification of mCherry/YFP fluorescence in the mutants with *GCG1*pr FPR background. The data includes values from three replicates and are normalized to the signal values of the wild type (FPR) strain. (D) Endogenous transcript analysis by RT-qPCR in the mutants. The bars represent fold gene expression ratio of coding and divergent transcript normalized to the expression of the reference gene, *UBC6*. Data information (C-D): The error bars represent SEM. Student’s t-test analysis shows the statistical significance of mutants compared to the respective wild type. *, **, *** and n.s. denote p-values <0.05, <0.01, <0.001 and non-significance, respectively.

### Hda1 and Hda3 repress endogenous DNC transcription

We selected the *GCG1/SUT098* locus for further validation of the top candidate regulators from the reporter screen. First, we performed an independent transformation of gene deletions into the *GCG1*pr FPR background to validate the effect. Quantification of mCherry/YFP fluorescence confirmed that the *hda1*Δ and *hda3*Δ mutants significantly reduced the fluorescence ratio, and the SAGA mutants increased the ratio (Figure 2C). These data are consistent with the data from the high-throughput reverse genetic screen, where Hda1C and SAGA scored on opposite sides. Second, to test the effect of the mutants on endogenous transcription, we quantified the expression of *GCG1* and *SUT098* transcripts by RT-qPCR. The *hda1*Δ and *hda3*Δ mutants significantly decreased the *GCG1/SUT098* ratio (Figure 2D) validating our findings. However, the SAGA mutants failed to increase the ratio at the endogenous level (Figure S2). We thus pursued the effects of Hda1 and Hda3 on DNC transcription.

We next asked whether the two subunits of Hda1C repress DNC transcription genome-wide. To address this question, we performed NET-seq in the WT, *hda1*Δ, and *hda3*Δ strains. To generate a systematic computational analysis, we first addressed the limited annotation of non-coding transcripts in the Saccharomyces Genome Database (SGD). We called novel DNC transcripts through an annotation algorithm incorporating NET-seq, Direct RNA-seq, CAGE-seq and 3’ READS data (See Materials & Methods). We identified 3736 novel non-coding transcripts in total, of which 1517 represent DNC transcripts that originate from the same NDR as the corresponding mRNA in the pair (Table S3). Among the identified DNC transcripts, 683 correspond to known CUTs or SUTs, whereas the other 834 were called *de novo*. Importantly, the metagene analysis of DNC loci revealed increased DNC levels in both *hda1*Δ and *hda3*Δ mutants compared to the WT signal (Figure 3A). These data are thus consistent with Hda1C as a genome-wide repressor of DNC. Since we identified the effect of Hda1C through *SUT098* expression, which represents a highly expressed DNC, we next analyzed if the Hda1C effect may be determined by the DNC expression level. To this end, we classified the DNC loci into three groups based on their level of nascent transcription. We found that Hda1C mutants increased DNC transcription irrespective of the expression levels (Figure S3A-C). Interestingly, the Hda1C mutants increased nascent DNC without a detectable effect in the direction of protein-coding transcription. We quantified the mutant and WT NET-seq signal at a locus with an annotated DNC transcript, *YPL172C/CUT409* (Figure 3B), and a locus with a DNC transcript identified through our analyses at *YDR261C* (Figure 3C) and detected increased DNC transcription at both loci in Hda1C mutants. The Hda1C mutants also increased DNC transcription at *GCG1* and *ORC2* (Figure S3D-E). In conclusion, the nascent transcriptome analyses supported the conclusion that Hda1 and Hda3 repress DNC transcripts genome-wide.

**Figure 3.**
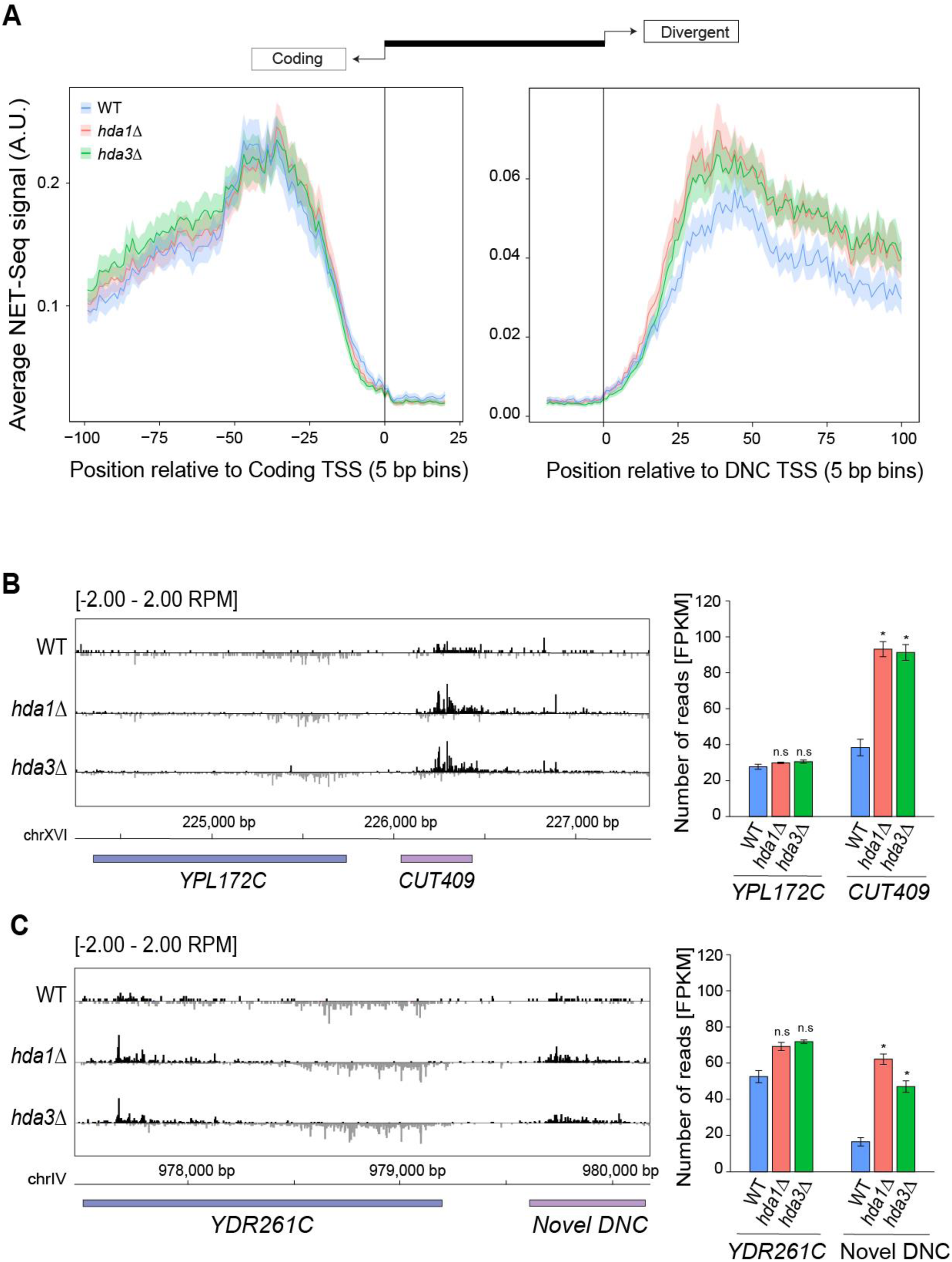
NET-seq identifies the genome-wide effect of Hda1C on nascent divergent non-coding (DNC) transcription. (A) Metagene analysis of NET-seq signal at all DNC loci (n=1517). The genomic intervals were centered at the transcription start site (TSS) of either protein-coding gene (left panel) or DNC transcript (right panel). (B) NET-seq data at the *YPL172C* locus. The signal represents NET-seq reads showing nascent transcription at the genomic position of the divergent non-coding strand (black) and the coding strand (grey). The bar graph depicts NET-seq reads as FPKM values for the coding and DNC transcript in the strains. Error bars represent SEM. Statistical significance of mutant transcript expression as compared to WT was assessed by student’s t-test. The * and n.s indicate p-value <0.05 and non-significance, respectively. (C) NET-seq data at the *YDR261C* locus with a novel DNC transcript. Annotations as in (B).

### Hda3 alters the frequency of DNC transcription at the GCG1/SUT098 locus

To investigate the molecular mechanisms of DNC repression by Hda1C we utilized the MS2/PP7 RNA-aptamer-based system to monitor live transcription at the *GCG1/SUT098* locus. We inserted stem-loop repeat sequences in the 5’ UTR of the endogenous genomic transcript sequences, 12xMS2 for *GCG1* and 14xPP7 for *SUT098* (Figure 4A). The orthogonal expression of the fluorescently tagged MS2-mScarlet and PP7-GFP coat proteins enabled the visualization of transcription in real-time as described previously (Lenstra & Larson, 2016). We recorded transcription over time for 130 cells on average to quantify the transcription parameters namely, duration and frequency of transcription initiation (Figure 4B). The live-cell imaging revealed rare transcription initiation events in the coding direction (i.e. *GCG1*), and higher expression for the divergent *SUT098* transcript (Figure 4C, S4). The relatively stronger transcription of *SUT098* compared to *GCG1* was consistent with the transcript counts in the nucleus and whole-cell by single-molecule fluorescent in-situ hybridization (smFISH) (Figure S4), and the DNC increase detected in NET-seq data (Figure 1A). The signals for *GCG1* and *SUT098* resembled Poisson distributions, supporting constitutive expression rather than transcriptional bursting (Figure S4). The constitutive expression of *SUT098* is consistent with measurements of the *GAL10* ncRNA that is also transcribed constitutively (Lenstra *et al*., 2015). The rare transcriptional initiation of *GCG1* precluded an in-depth analysis of transcription kinetics. Although higher than *GCG1*, the transcriptional activity of *SUT098* is still low, which may also explain the lack of significant increase of *SUT098* transcript in the mutants by smFISH analysis. We thus restricted our studies on the quantification of the *SUT098* live-cell imaging data for the characterization of mutants. We compared the transcription duration of transcription in WT, the Hda1C mutants and the CAF-I mutant *cac2Δ* (Figure 4D). Our analyses identified no difference in transcription duration between the mutants and WT. Since these are individual transcripts, a similar transcription duration indicates that Hda1C does not affect the transcription elongation rate, but perhaps the initiation rate. Indeed, the *hda3*Δ mutant significantly decreased the time between transcription events, thus increasing the frequency of DNC transcription initiation (Figure 4D). Although not statistically significant, we note that the data indicated the same trend towards an increase of DNC transcription frequency also in *hda1Δ* and *cac2Δ*. In summary, the live-cell imaging data suggested that increased DNC in the *hda3*Δ mutant may result from increased transcriptional initiation frequency.

**Figure 4.**
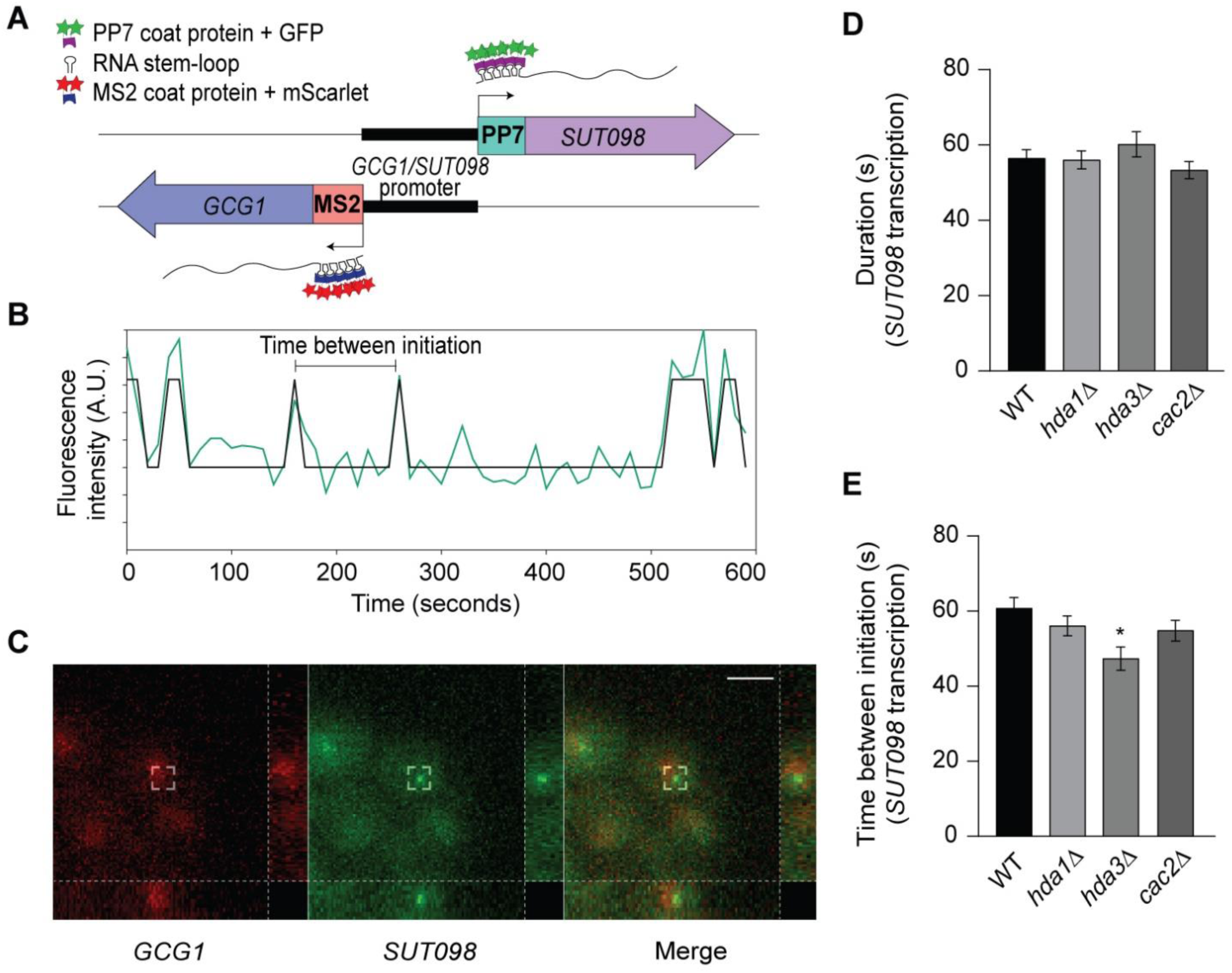
Single-molecule live-cell imaging reveals *hda3*Δ altering *SUT098* transcription frequency. (A) Schematic representation of MS2/PP7 stem-loop repeats at the 5’UTR of *GCG1/SUT098* sequence, respectively. Upon transcription, respective coat proteins (dark blue and dark purple) with fluorescent proteins (red and green) bind to the MS2/PP7 repeats at the 5’ UTR of *GCG1* and *SUT098* transcript. The ON and OFF state of transcription versus the time determines the duration and frequency (time between transcription events) of transcription. (B) An example trace of *SUT098* transcript fluorescence quantified at the transcription site in a single cell (wild type). The track shows an overlay of the raw trace (green) with the binarized trace (black). The binarized peaks represent transcription initiation. (C) A representative image of transcription initiation observed as fluorescence spots in the recorded live-cell movie of wild type strain. The red and green channel denotes *GCG1* and *SUT098* transcript initiation, respectively. Images represent maximum intensity projections, and the right and bottom sidebars indicate sideviews in the yz and xz directions, respectively. Scale: 5 µm. (D, E) Quantification of transcription dynamics by the duration and frequency (time between initiation) of transcription. Bar graphs show the transcription parameters of *SUT098* in candidate mutants compared to the wild type strain. The error bars show SEM. Statistical significance was calculated by bootstrapping (the asterisk * denotes p-value <0.05).

### Hda1 affects H3 acetylation at DNC loci and acts independently of the H3K56 pathway

We next focused on links between histone acetylation and the repression of DNC transcription by Hda1C. The H3K56ac pathway contributes to DNC repression (Marquardt *et al*., 2014). We performed a genetic epistasis analysis to test if the effect of Hda1C on DNC could be explained through H3K56ac. Point mutations in H3K56 to A or Q affect DNC transcription (Marquardt *et al*., 2014; Rege *et al*., 2015). If the effects of Hda1C were explained by H3K56ac, we would expect the same effect on DNC transcription in H3K56/*hda* double mutants as in the single mutants. We detected an increased ratio of mCherry/YFP fluorescence of the *GCG1*pr FPR in K56Q, K56A, and H3WT/*hda1*Δ (Figure 5A). Importantly, the fluorescence ratio was further increased in H3 K56Q/*hda1*Δ and K56A/*hda1*Δ double mutant combinations compared to the single mutants. These data thus revealed an additive effect of the H3K56ac and *hda1*Δ pathways on DNC transcription, suggesting that H3K56ac and Hda1C make non-overlapping contributions to DNC repression. This may be explained by the fact that Hda1C does not only deacetylate H3K56, but also other residues at H2B, H3 (Carmen *et al*., 1996), as well as H4 at active genes (Ha *et al*., 2019). We therefore investigated H3 acetylation levels at DNC loci genome-wide with ChIP-seq in WT and *hda1*Δ (Ha *et al*., 2019). H3 acetylation signal increased in the DNC direction in *hda1*Δ compared to WT (Figure 5B). These data reinforce the view that increased transcription frequency and elevated nascent DNC transcription in the Hda1C mutants are linked to elevated histone acetylation.

**Figure 5.**
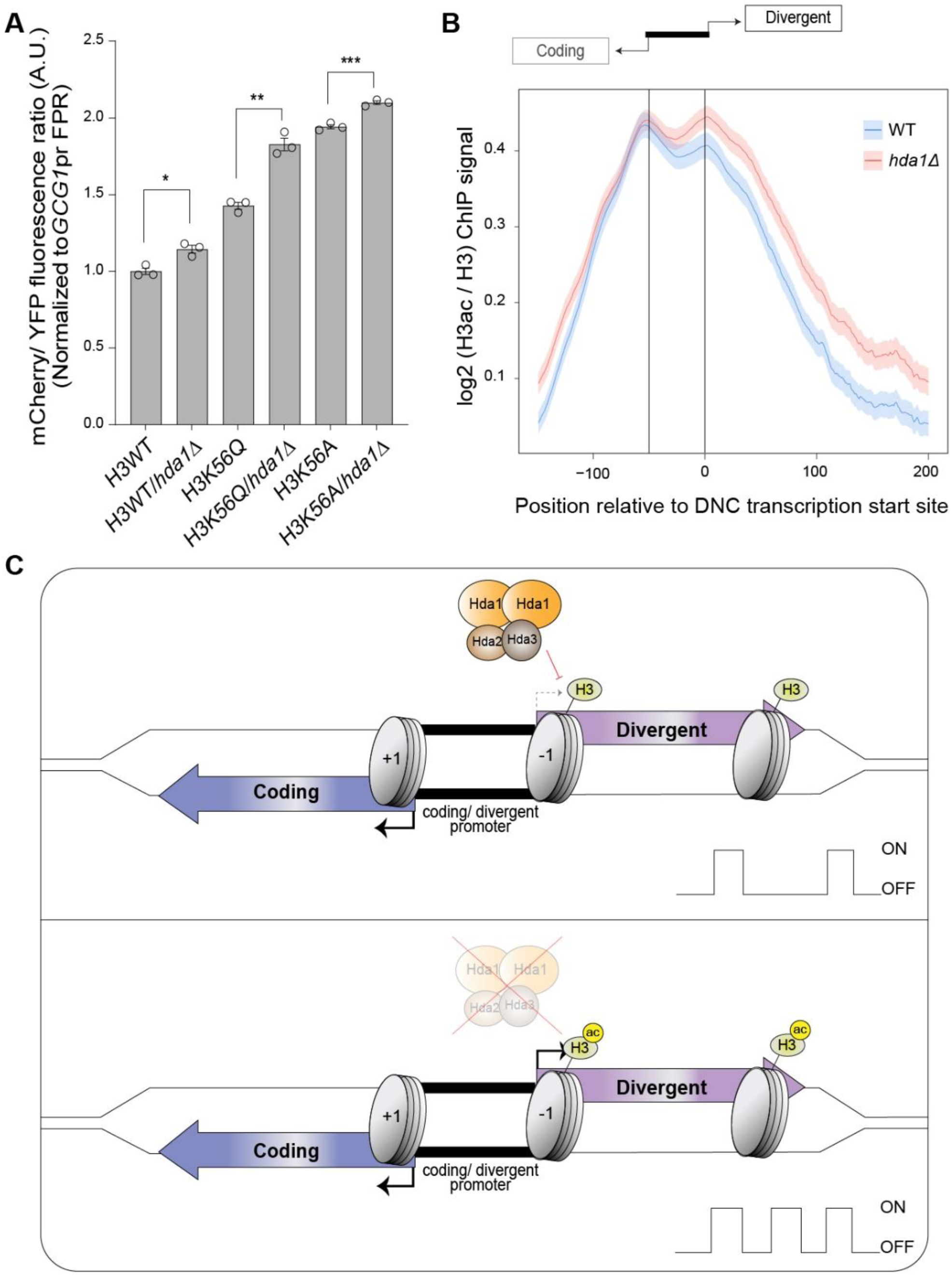
Hda1 affects the H3 acetylation levels at divergent non-coding (DNC) region independently from H3K56ac. (A) Flow cytometry data analysis. Bars represent the mCherry/YFP fluorescence ratio of the respective yeast mutants normalized to the signal values of the H3 wild type FPR strain. The error bars show SEM. Student’s t-test analysis indicates the statistical significance of the mutant compared to the respective wild type. The asterisks *, ** and *** indicate p-values <0.05, <0.01 and <0.001, respectively. (B) Metagene profile of ChIP data from (Ha *et al*., 2019). The genomic intervals cover the first 500 bp of the coding gene (scaled to 100 bins), the first 1 kb of the DNC transcript (scaled to 200 bins), and the variable length gap between the coding TSS and the DNC TSS (scaled to 50 bins). The H3ac ChIP-seq signal was normalized by the H3 signal in the same samples. (C) Working model of Hda1C repressing DNC. The coding (blue) and divergent (purple) transcript arise in opposing directions of a shared nucleosome-free region (black). Nucleosomes (grey) comprise histones capable of modification by histone deacetylase 1 complex (Hda1C). In the normal state, Hda1C limits the frequency of DNC by deacetylating histones. With the loss of Hda1C function, increased acetylation of H3 favors DNC.

## DISCUSSION

Previously, we identified a key contribution to H3K56ac-mediated histone exchange in DNC transcription based on a genetic screen with the *PPT1/SUT129* regulatory region. The incorporation of H3K56ac nucleosomes by the histone chaperone CAF-I restricted DNC transcription (Marquardt *et al*., 2014). Two main factors motivated new genetic screens: i) we failed to resolve protein complexes necessary for DNC (i.e. activators), and ii) our genome-wide characterization of NET-seq data in *cac2Δ* mutants suggested DNC regulation independent of the H3K56ac pathway.

We reasoned that loci with high DNC transcription levels should facilitate the detection of mutations reducing DNC transcription. Replacement of endogenous transcripts with mCherry/YFP offers the advantage of screening for regulation upstream of ncRNA repression by co-transcriptional RNA degradation (Wyers *et al*., 2005; Neil *et al*., 2009; Van Dijk *et al*., 2011; Ntini *et al*., 2013), but some hits may be specific to the FPR. Consistently, we did not identify genes regulating DNC through transcriptional termination and co-transcriptional RNA degradation. Our genetic screen resolved protein complexes as activators, for example, SAGA, ISW2, and confirmed the role of SWI/SNF (Marquardt *et al*., 2014). The HAT activity of SAGA appeared particularly promising in light of the HDACs identified as repressors. However, we failed to validate reduced DNC transcription for the endogenous transcripts in activator mutants. While genome-wide methods could possibly reveal a reduction of DNC transcription, we consider it likely that our screen failed to identify genuine activators of DNC. In addition to FPR-specific effects, it is more likely that SAGA, SWI/SNF and ISW2 protein complexes regulate the transcription of mRNAs rather than DNCs given previous reports (Hassan *et al*., 2001; Whitehouse *et al*., 2007; Baptista *et al*., 2017; Kubik *et al*., 2019), arguing against an unknown complex dedicated to promote DNC transcription specifically. Nevertheless, since we screened against a library of non-essential gene deletion mutants, it remains possible that a protein complex essential for yeast viability could stimulate DNC transcription. Our computational analyses suggest DNC at about 28% of yeast genes, with 1517 DNC transcripts compared to 5544 expressed yeast genes (i.e. meta-NET-seq data, FPKM above 10). Even though DNC is frequent, yeast genes without evidence for DNC are thus in the majority. One explanation for a large number of genes without DNC may be reduced activities of elusive DNC activators. Alternatively, DNC transcription may result from NDR formation linked to mRNA expression as suggested by the transcriptional noise hypothesis (Struhl, 2007). However, locus-specific repressor activity reduces DNC transcription, resulting in variations of DNC transcription genome-wide. Our data thus suggest to extend the transcriptional noise hypothesis with activities limiting DNC transcription to account for genome-wide variation in non-coding transcription.

Our screens highlighted the role of Hda1C as a repressor of DNC transcription. In budding yeast, Hda1C comprises the Hda1, Hda2, and Hda3 subunits and has a distinct function from other HDACs (Wu *et al*., 2001). The loss of the catalytic subunit Hda1 or either of the two functional subunits disrupts the activity of the complex (Lee *et al*., 2009). We note that our analyses failed to identify the equivalent effects in the respective mutants, perhaps most notably a statistically significant increase of initiation frequency was limited to *hda3*Δ. While the differences may be rooted in differential effects of the Hda1C components (Lee *et al*., 2021), we favor the hypothesis that some effects may be masked by experimental variation and the modest effect size. Hda1C selectively deacetylates histones H2B and H3 (Carmen *et al*., 1996). The acetylation states at lysine residues of histones are linked to transcription dynamics which in turn control the gene expression (Wu *et al*., 2017). Hda1C activity is genetically separable from the H3K56ac-linked DNC repression mediated by histone chaperones that involve the Hst3/4 histone deacetylase acting on H3K56 (Marquardt *et al*., 2014). We detected clear histone acetylation signals at DNC loci (Figure 5B), consistent with the idea that HDACs may modulate DNC transcription (Churchman & Weissman, 2011; Tan-Wong *et al*., 2012). The additive genetic interaction between H3K56 and *hda* mutants supports a parallel contribution of both H3K56ac and Hda1C mediated mechanisms to limit DNC transcription. We thus propose a model highlighting DNC repression by Hda1C through counteracting histone acetylation beyond H3K56ac (Figure 5C). In conclusion, histone acetylation emerges as a key chromatin-based facilitator of DNC transcription.

In order to address the mechanism of DNC repression, we implemented live-cell imaging. Since open chromatin at promoters is linked to histone acetylation (Barnes *et al*., 2019), the acetylation state may favor initiation from promoters (Dar *et al*., 2012). We identified a reduction in time between DNC transcription initiation events when Hda1C-mediated histone deacetylation is impaired. Our data support a model where HDACs repress DNC transcription through a reduction in the frequency of transcription initiation (Figure 5C). We note that increased histone acetylation increases the transcription frequency also in mouse (Chen *et al*., 2019), perhaps indicating that histone acetylation may correlate with increased transcription frequency generally. Unfortunately, the rare *GCG1* transcription initiation events at the *GCG1/SUT098* promoter precluded a simultaneous comparative analysis of the effects of histone de-acetylation in both transcriptional directions. Nevertheless, our results suggest that histone deacetylation by Hda1C limits DNC transcription through a reduction in transcriptional initiation frequency.

In conclusion, even though DNC transcription is frequent and occurs at high levels in the yeast genome (Xu *et al*., 2009), HDACs reduce DNC initiation frequency to maintain tolerable levels of pervasive transcription. Presumably, the advantages of harnessing the contributions of selected DNC transcription events for gene regulation outweigh the penalties of widespread DNC activity. Since no dedicated factors stimulating genome-wide DNC could be identified thus far, we favor the idea that DNC transcription follows from promoter NDRs without the need for specific pathways activating DNC. The differences in DNC levels across yeast promoter NDRs thus likely result from DNC regulation through a series of parallel pathways for repression, including the control of initiation frequency by histone deacetylation, and through co-transcriptional ncRNA degradation.

## Supporting information

Description of supplementary files

Table S4 yeast strains

Table S5 oligonucleotides

Table S6 plasmids

## Supplementary Figures

**Figure S1.**
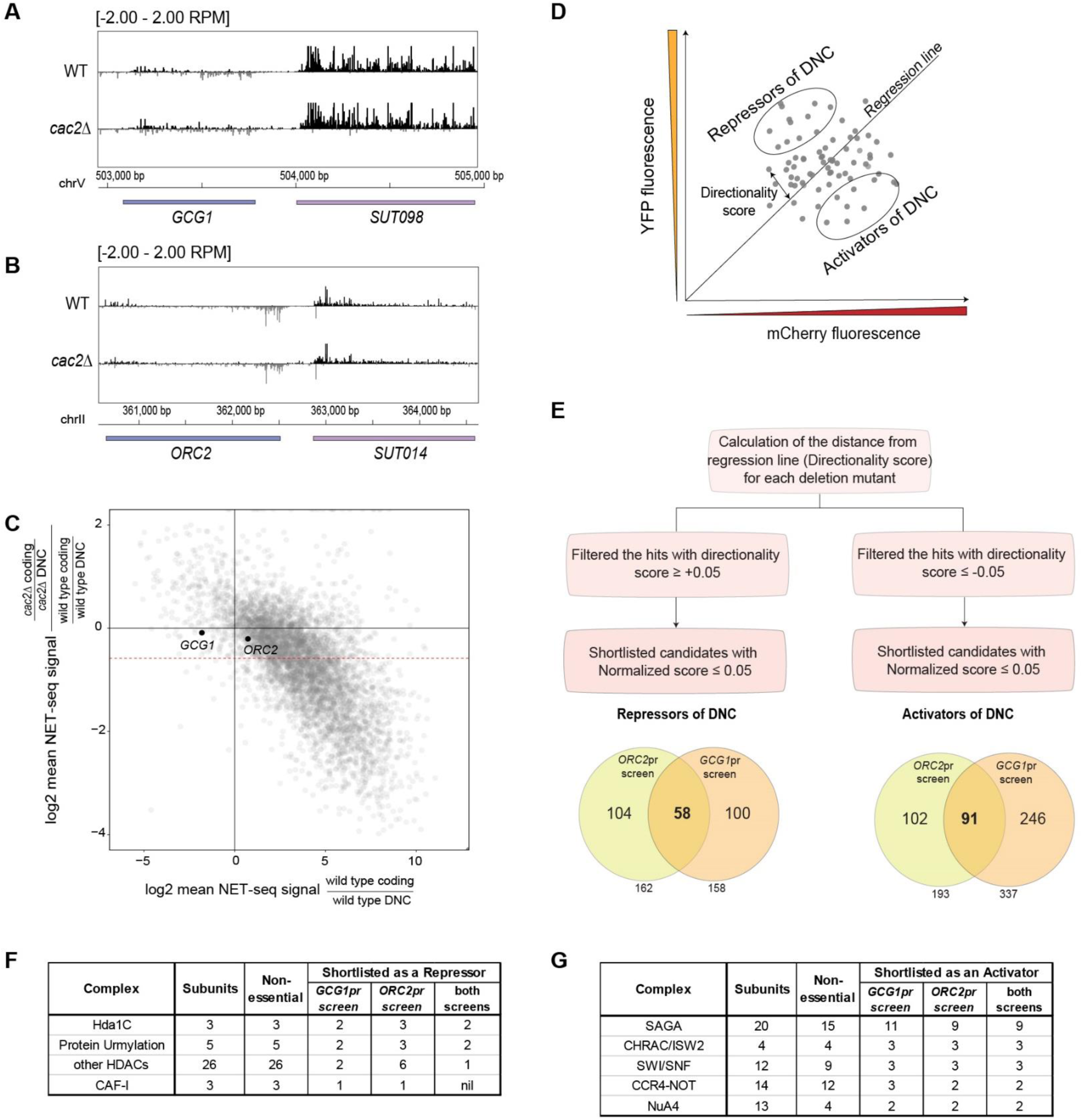
Selection of loci to find factors regulating divergent non-coding (DNC) transcription and method of reporter screen data analysis. (A, B) NET-seq data from Marquardt *et al*., 2014 at the *GCG1/SUT098* and *ORC2/SUT014* genomic loci. The two tracks represent normalized signal values in WT and *cac2*Δ. The black and grey signals distinguish the Watson and Crick strand, respectively. (C) NET-seq coding/non-coding ratios in *cac2*Δ and WT. For each gene, the directionality was calculated as the ratio of coding to divergent non-coding reads. The CAF-I effect was measured as the ratio of directionality in the mutant divided by that in WT and plotted against the WT directionality using a log2 scale. Changes in the mutant appear as deviations from the zero value on the y-axis. A 1.5-fold cutoff marking CAF-I affected loci is marked by a dashed red line. Highlighted in black are the *GCG1* and *ORC2* promoters unaffected by *cac2*Δ. (D) Schematic illustration of data from a reporter screen. Plotting the median YFP versus mCherry fluorescence signal results in a scattered distribution of data points each representing a deletion mutant. The regression line distinguishes mutants favoring divergent or coding transcription. The directionality score is proportional to the distance from each data point to the regression line. Mutants favoring DNC transcription (i.e. with positive directionality values) are the Repressors of DNC. Mutants favoring coding transcription (found below the regression line) are the Activators of DNC. (E) Depiction of flow cytometry data analysis. Calculation of directionality scores from individual reporter screens allowed in the classification of mutants as repressors and activators of DNC (Table S2). The Venn diagram represents the number of shortlisted candidate repressors and activators in each screen. (F, G) List of protein complexes short-listed in the genetic screens. The table summarizes the number of sub-units of the complexes and the non-essential sub-units found in the individual genetic screens.

**Figure S2.**
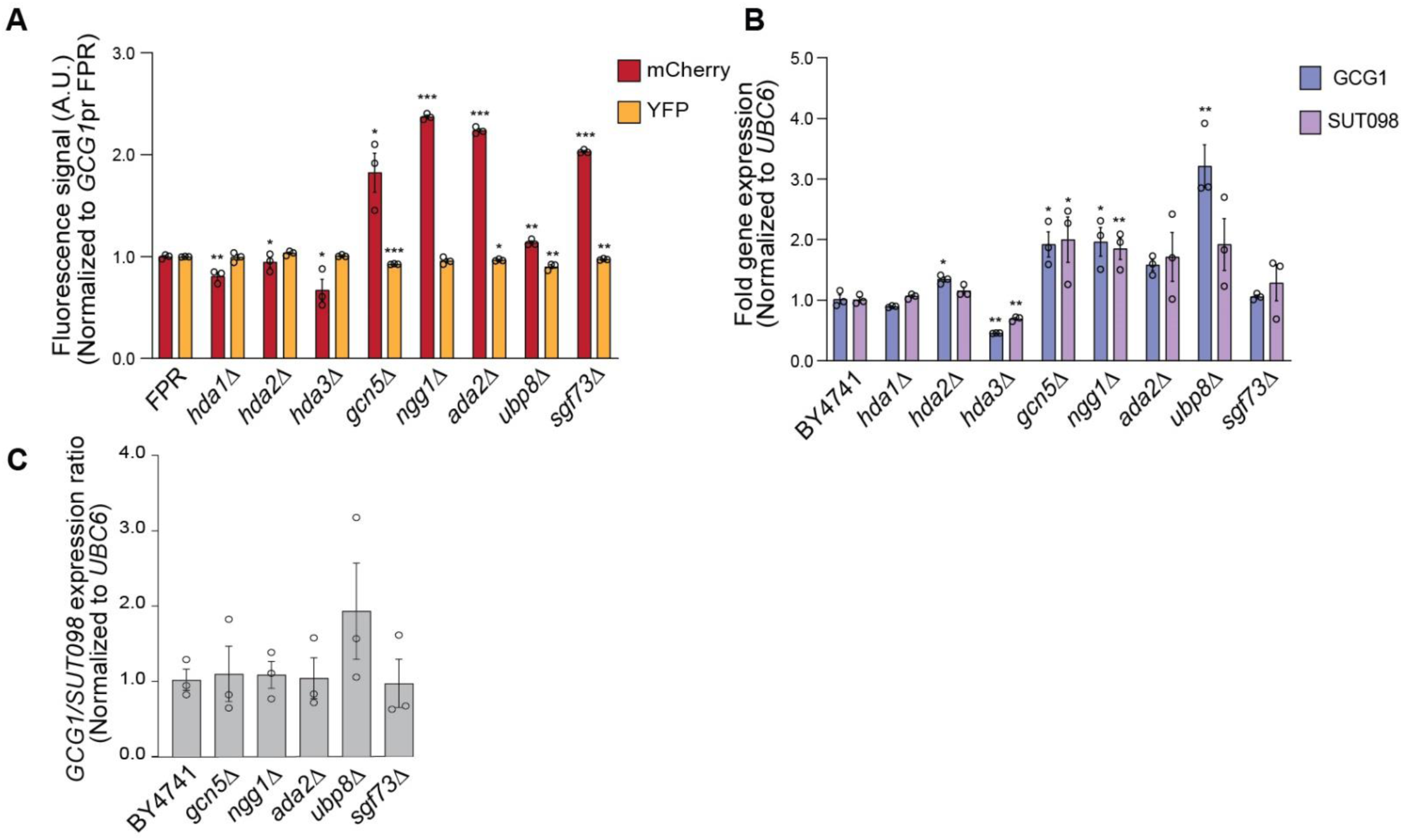
Quantification of coding and divergent non-coding (DNC) transcripts in candidate mutants. (A) The fluorescent mCherry and YFP protein signals quantified by flow cytometry data analysis with the *GCG1*pr FPR construct. The individual signal values were normalized to the FPR signal values. (B) Endogenous levels of the coding *GCG1* and the divergent *SUT098* transcript as measured by RT-qPCR. The fold gene expression values were normalized with the *UBC6* reference gene expression. (C) Ratio of endogenous coding versus the DNC transcript fold gene expression at the *GCG1*/*SUT098* locus. Data information (A-C): The data includes values from at least three replicates and the error bars represent SEM. Student’s t-test analysis shows the statistical significance of mutants compared to the respective wild type. The asterisks *, ** and *** represent p-values <0.05, <0.01 and <0.001, respectively.

**Figure S3.**
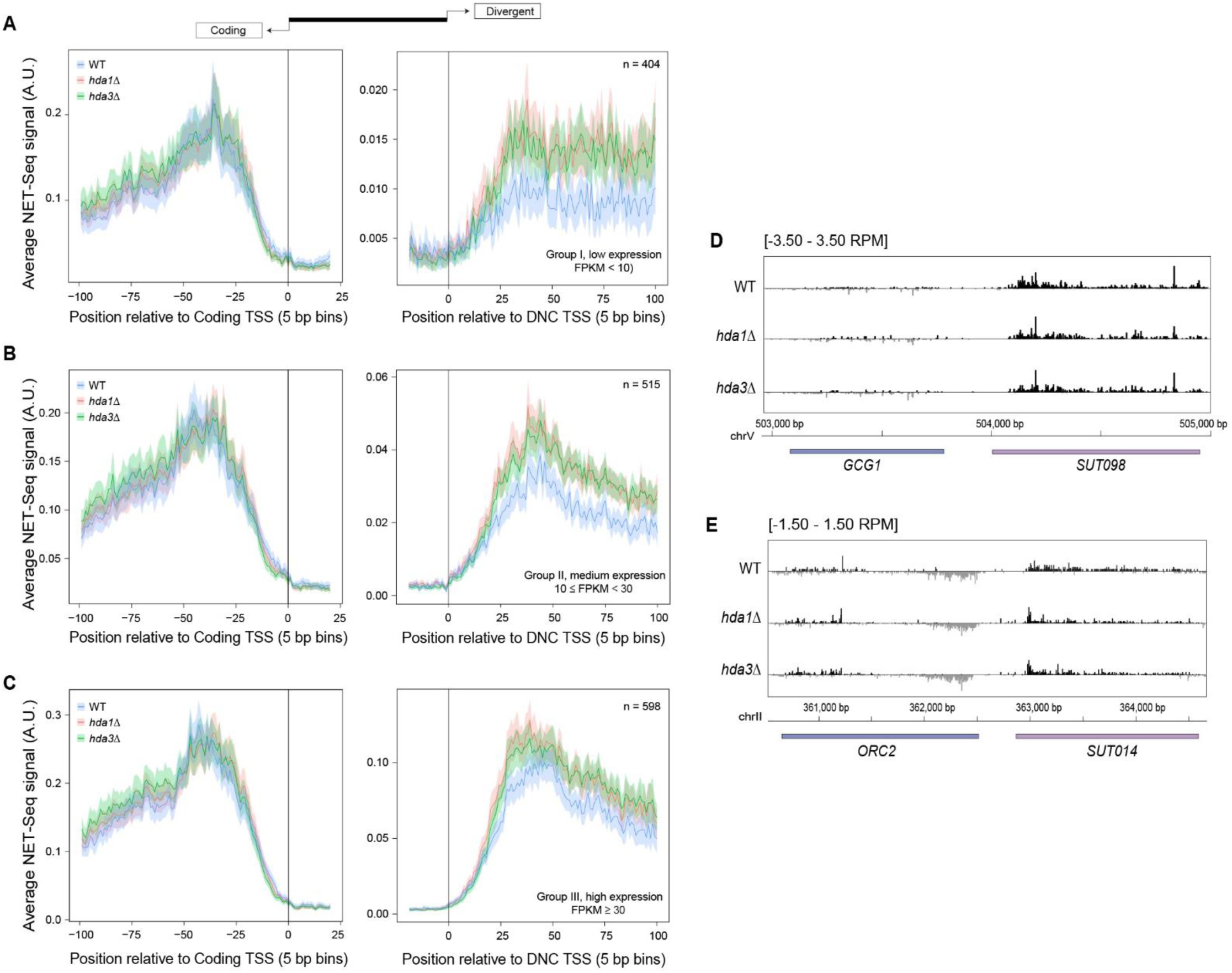
NET-seq data supporting the Hda1C effect on global nascent DNC transcription as shown in Figure 3. (A-C) Metagene analysis of NET-seq signal in wild type (WT), *hda1*Δ, and *hda3*Δ yeast strains stratified by the DNC expression level. The genomic intervals were centered at the TSS of the coding gene (left panel) and the DNC transcript (right panel). (A) Loci with low DNC expression (FPKM <10), n = 404. (B) Loci with medium DNC expression (10 ≤ FPKM < 30), n = 515. (C) Loci with high DNC expression (FPKM ≥ 30), n = 598. (D, E) NET-seq data track at the *GCG1*/*SUT098* and *ORC2/SUT014* loci. The tracks represent normalized signal values in WT, *hda1*Δ and *hda3*Δ. The black and grey signals distinguish the Watson and Crick strand, respectively.

**Figure S4.**
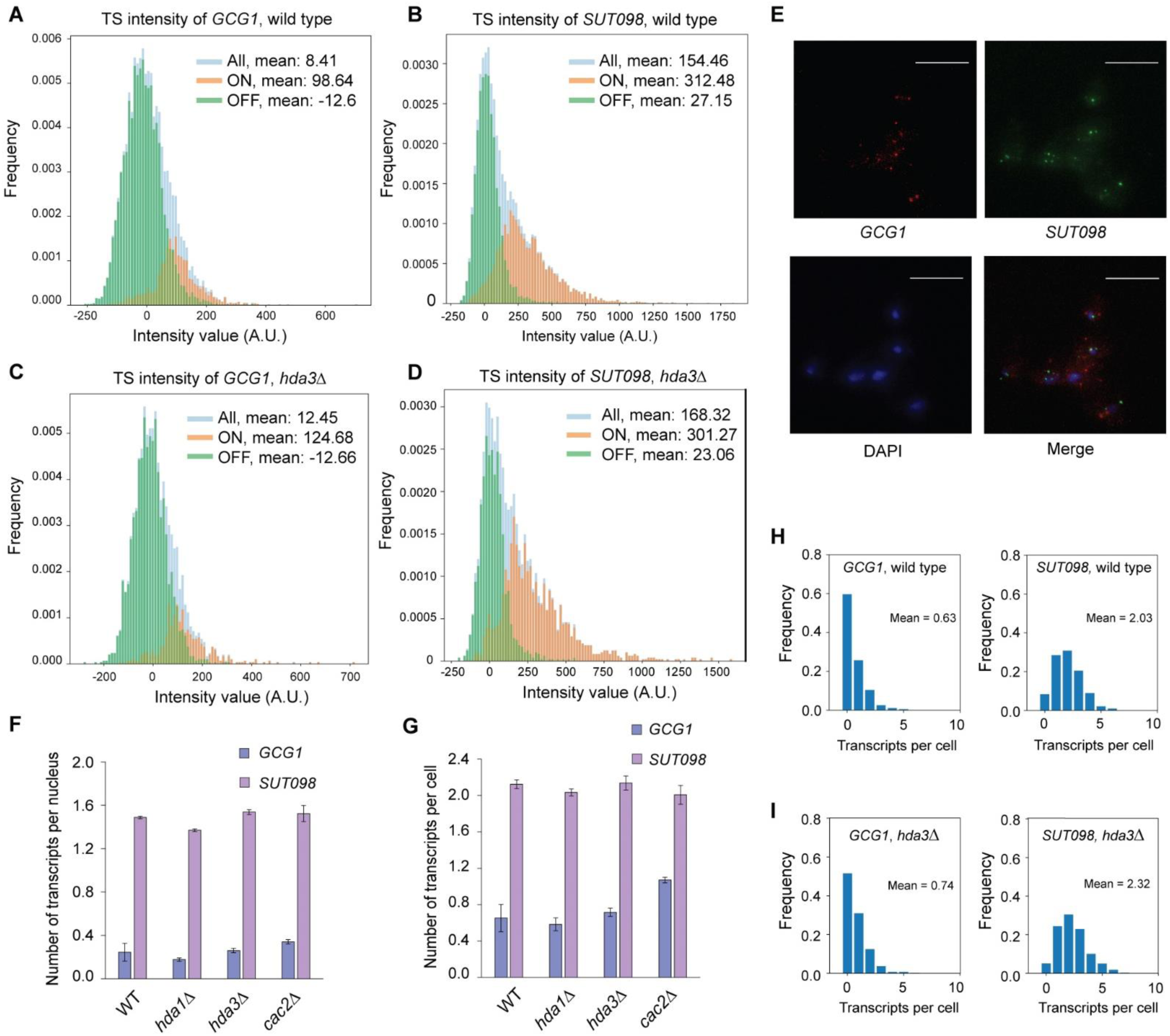
Live-cell imaging data analysis and quantification of transcripts at the *GCG1/SUT098* locus by single-molecule fluorescence in-situ hybridization (smFISH). (A, B) Intensity of *GCG1* and *SUT098* transcript at the transcription site (TS) by live-cell imaging in wild type cells. The bars represent fluorescence intensity distribution of all the analyzed single cells (light blue), cells during initiation (ON, orange) and cells after initiation (OFF, green). (C, D) Intensity of *GCG1* and *SUT098* transcripts at the transcription site (TS) by live-cell imaging in *hda3*Δ cells. The bars represent fluorescence intensity distribution of all the analyzed single cells (light blue), cells during initiation (ON, orange) and cells after initiation (OFF, green). (E) Representative image of the *GCG1* and *SUT098* transcripts in wild type strain, by smFISH. The channels show wide-field images of fixed haploid cells. The blue is DAPI, red and green dots are the *GCG1* and *SUT098* transcripts, respectively. The scale bar is 10 µm. (F) Average count of the *GCG1* and *SUT098* transcripts in the nucleus by smFISH. (G) Whole-cell count of the *GCG1* and *SUT098* transcripts by smFISH. (H, I) Representative histogram of the number of transcripts per cell in a wild type (n=1409) and *hda3*Δ (n=1593) replicate. The bars indicate the frequency of the transcripts per cell. The mean transcript number and the number of cells analyzed are indicated inside the panels. Data information (F, G): The data contains at least three replicate values. The data analysis and representation include the cells without any transcripts to account for the loss of transcription initiation. The error bars represent SEM.

**Figure S5.**
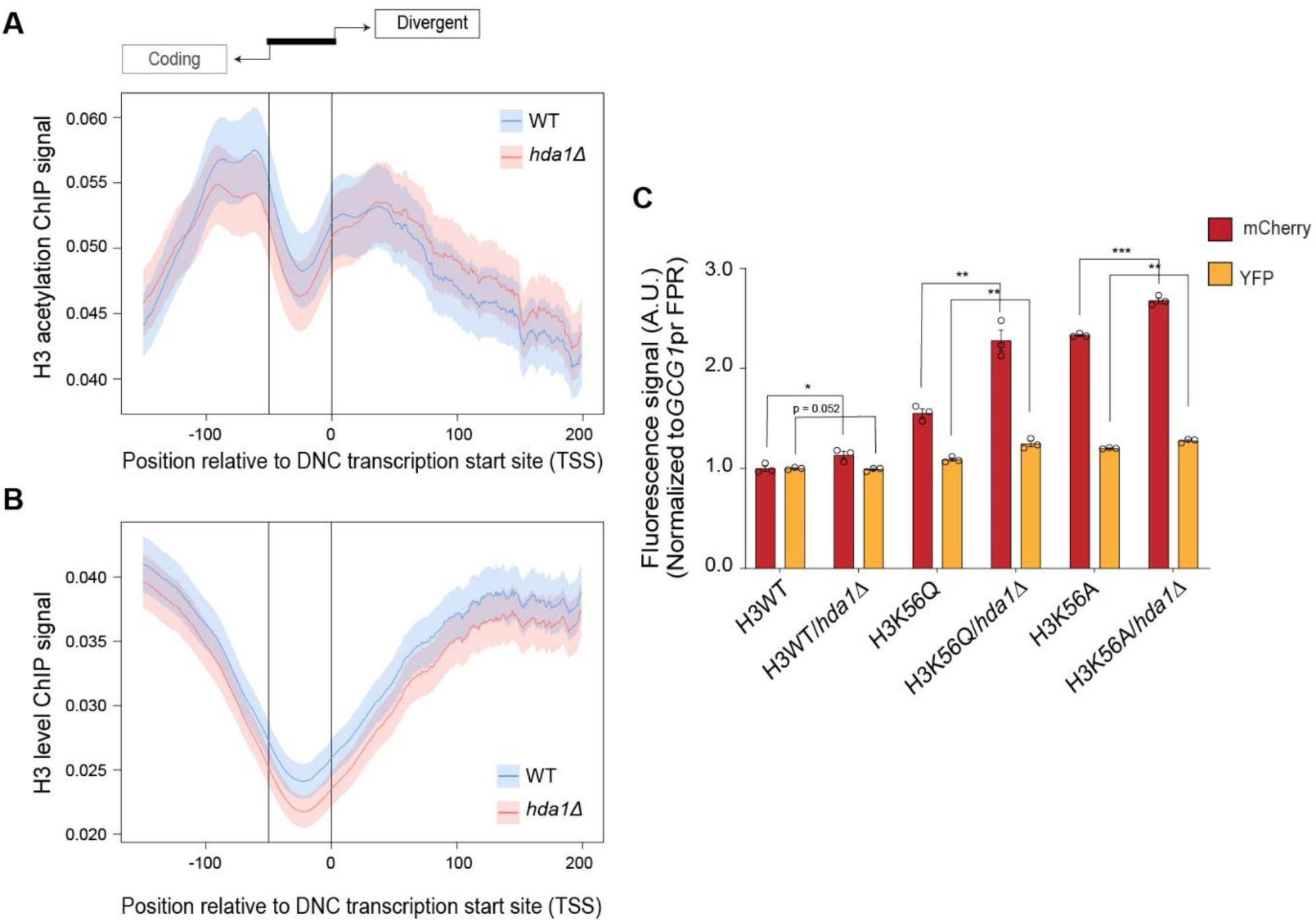
Hda1C deacetylates H3 in divergent non-coding (DNC) regions. (A, B) Metagene profiles of H3 acetylation and H3 ChIP signal from (Ha *et al*., 2019) at DNC loci. The signal represents the (A) acetylation levels of H3 and (B) H3 levels. The genomic intervals cover the first 500 bp of the coding gene (scaled to 100 bins), the first 1 kb of the DNC transcript (scaled to 200 bins), and the variable length gap between the coding TSS and the DNC TSS (scaled to 50 bins). (C) The fluorescent mCherry and YFP protein signals quantified by flow cytometry data analysis in H3 WT background strain with the *GCG1*pr FPR construct. The individual signal values are normalized to the FPR signal values. The data includes values from at least three replicates and the error bars represent SEM. Student’s t-test analysis shows the statistical significance of mutants compared to the respective wild type. The asterisks *, ** and *** represent p-values <0.05, <0.01 and <0.001, respectively.

## MATERIAL & METHODS

### Growth media

#### Yeast (*S. cerevisiae*)

Standard YPD liquid media or agar plates with 500 µg/ml G418 sulfate (Abcam) and/or 100 µg/ml clonNAT (Werner BioAgents) and/or 200 µg/ml Hygromycin B (Invitrogen) were used to select for yeast strains with KanMX or NatMX or HphMX resistance, respectively. Synthetic complete drop-out media (minus URA/ HIS/ LEU) was used to select for strains with an auxotrophic marker gene.

#### Bacteria (*E. coli*)

Standard LB liquid or agar plates with 100 µg/ml Ampicillin or 50 µg/ml Kanamycin were used to select cells harboring the plasmid vector with Amp^R^ or Kan^R^ selection marker.

### Cloning

#### *GCG1*pr and *ORC2*pr FPR constructs

The *PPT1/SUT129* promoter present in SMC50 (Marquardt *et al*., 2014) was replaced with the bidirectional *GCG1/SUT098* or *ORC2/SUT014* promoter sequence, respectively using isothermal assembly reaction (Gibson *et al*., 2009). The FPR cassette comprising NatMX-mCherry-*PPT1/SUT129* promoter-YFP was excised from the backbone using AscI restriction enzyme (1 U per µg plasmid, NEB). Fragment I was the bidirectional *GCG1/SUT098* or *ORC2/SUT014* promoter sequences amplified by Phusion U polymerase from BY4741 genomic DNA. Fragment II and III were the YFP and mCherry sequences with complementary overhangs to the respective promoters (introduced by primer overhangs corresponding to about 20 basepairs of the bidirectional promoter) amplified from SMC50 plasmid. The three fragments were cleaned up using the Wizard^®^ SV Gel and PCR Clean-Up System (Promega) and subsequently fused by overlapping PCR. The fused product was ligated into the SMC50 vector backbone by the isothermal assembly reaction. The ligated plasmid was transformed into *E. coli* and sequenced. For transformation into yeast, the plasmid can be linearized with EcoRI and the plasmid integrates into the *PPT1/SUT129* locus of the genome.

#### *GCG1*pr FPR cassette with KanMX resistance

The *GCG1/SUT098* FPR plasmid was digested with AvrII and BglII (1 U per µg plasmid, NEB) restriction enzymes to excise the NatMX6 cassette. A KanMX6 cassette with overhangs complementary to the opened site of the backbone was amplified using Phusion U polymerase chain reaction and fused into the backbone by an isothermal assembly reaction.

### Transformation

Yeast transformations were performed as described in (Gietz & Schiestl, 2007) with minor changes. For each transformation, cell pellet collected from 5 ml of 0.5-0.8 OD_600_ liquid culture was washed 2x in sterile water and 1x in sterile 100 mM Lithium Acetate. The cells were resuspended in 74 µl of 1 - 2 µg of PCR amplicon or 100 - 500 ng of plasmid or 1 µg of the linearized plasmid. A sterile transformation mix consisting of 240 µl 50% w/v PEG 3350, 36 µl 1 M Lithium acetate, and 10 µl sheared salmon sperm DNA (10 mg/ml) was added to the tube. The contents were vortexed briefly and incubated for 30 min, RT (with frequent mixing), and heat-shocked for 20 min, 42°C in a heat block. The transformed cells were collected and washed 1x with sterile water. For selection with auxotrophic marker, the transformed cells were plated directly on synthetic drop-out media. For selection with antibiotic markers, the cells were plated on plain YPD and replica plated onto selection media the next day or plated directly on selection media after 3-4 hours of recovery in 1 ml of fresh YPD at 30°C, 200 rpm.

### Synthetic Genetic Array (SGA)

Query strains (MATα) harboring *ORC2*pr FPR construct (SMY2312) or *GCG1*pr FPR (SMY2314) were crossed with the yeast non-essential gene deletion library (MATa) as described in (Baryshnikova *et al*., 2010) to perform the reverse genetic screen. The resultant MATa haploid cells from SGA on 384-well plates were quantified for fluorescence in the flow cytometer.

### Flow cytometry

Quantification of mCherry and YFP fluorescent signal was done using an LSR II flow cytometer from Becton Dickinson with a high-throughput sampler. YFP was excited at 488 nm and the emission was collected through a 545/35 band pass and 525 long pass emission filter. The mCherry was excited at 594 nm and the fluorescence was collected through a 632/22 band pass filter. 10000 - 50000 events were recorded for each well. The BD FACSDiva acquisition program was used to set up the flow settings and the acquired data were exported in FCS 3.0 format.

### RNA isolation

The total RNA from yeast cells was extracted using the phenol/chloroform method. Cells from 40 ml of 0.5-0.8 OD_600_ culture were collected, washed in 2 ml of sterile water, and resuspended in 400 µl of ice-cold AE buffer (50 mM sodium acetate, pH 5.0 and 10 mM EDTA, pH 8.0). 50 µl of 10% SDS and 500 µl of fresh phenol/chloroform mixture (1:1) were added to the tubes, vortexed for 5 min with glass beads, and incubated at 65°C, 800 rpm for 10 min. The tubes were chilled on ice for 5 min and centrifuged at 4°C, high speed to separate the contents. The aqueous phase was collected and mixed with 500 µl of fresh phenol/chloroform mixture (1:1), vortexed for 10 min, and incubated at 65°C, 2000 rpm for 5 min. The tubes were chilled on ice for 10 min and centrifuged at 4°C, high speed to separate the contents. The aqueous phase was transferred to fresh tubes, mixed with 2.5 volume of cold 100% ethanol, 1/10 volume of 3M sodium acetate, pH 5.3, and incubated at −20°C for 2 hours. The tubes were centrifuged at 4°C to pellet the RNA. The RNA pellet was washed with 75% ethanol and air-dried until complete removal of ethanol and re-suspended in 400 µl of RNase-free water. The isolated RNA was quantified using a nanodrop.

### qPCR

Total RNA from yeast cells was extracted using the standard hot phenol/chloroform method. The total RNA was treated with TURBO DNase (ThermoFisher Scientific) following the protocol from the manufacturer. 1 µg of the DNase treated RNA was converted to cDNA with Superscript IV Reverse transcriptase kit (Invitrogen, USA) using gene-specific primers based on the manufacturer’s instructions. Diluted cDNA (1:20) was used in a PCR reaction with the GoTaq qPCR Master Mix (Promega) and run on a CFX384 Touch instrument (Bio-Rad).

### qPCR analysis

The output data were exported to Microsoft Excel and analyzed. Quantification of the transcript expression relative to the *UBC6* internal reference gene was performed using the ΔΔCt method as described in (Livak & Schmittgen, 2001).

### Native Elongating Transcript Sequencing (NET-seq)

Nascent RNA was immune-precipitated from the wild type strain and the *hda1Δ* and *hda3Δ* mutants (two biological replicates for each genotype) using the previously published protocol (Churchman & Weissman, 2011). The NET-seq libraries were constructed using the Bioo Scientific Small RNA-seq kit v3 and the custom protocol (Kindgren *et al*., 2020).

#### Harvesting and grinding of cells

One liter of mid-log phase yeast cells were collected using vacuum filtration and immediately frozen in liquid nitrogen. All equipment used in handling the cells were pre-cooled using liquid Nitrogen. The frozen cell mass was powdered by grinding 10x at 15 Hz, 3 min in a Retsch Mixer mill.

#### Quality check and Sequencing

The constructed libraries were validated using High sensitivity DNA kit (Agilent Technologies) in a Bioanalyzer as per the manufacturer’s instructions. The libraries were sequenced on Illumina NovaSeq 6000 in PE150 mode at Novogene (https://en.novogene.com/). 39-82 million reads were obtained for each of the sequenced libraries.

### Data analysis

All supporting code was deposited at https://github.com/Maxim-Ivanov/Gowthaman_et_al_2021 and https://github.com/Uthra-Gowthaman/Gowthaman-et-al_2021.

The FCS files were processed using the 05-Load_flow_cytometry_data.R script which is based on the flowCore library (Hahne *et al*., 2009). Only wells with at least 100 cells were considered valid. The forward scatter (FSC) and side scatter (SSC) values were used to filter out cell aggregates and odd-shaped cells. For each valid well, the median YFP and mCherry fluorescence values were calculated. The directionality scores were calculated as the distance between the position of a well in two-dimensional space and the regression line obtained from all wells on a given 384-well plate. Scatterplots of the YFP and mCherry fluorescence values, as well as the directionality scores, were obtained using the ggplot2 library (see the 06-Scatterplots.R script).

The raw FASTQ files from the previously published NET-seq studies and the current study were aligned to the yeast genome using the 01-Remapping_published_NET-seq_datasets.sh and 02-Alignment_of_novel_yeast_NET-seq_data.sh scripts, respectively. The ChIP-seq BigWig files from the Ha *et al*., 2019 study were downloaded from NCBI GEO (accession number GSE121761). The NET-seq and ChIP-seq metagene plots were produced by the 07-Metagene_plots.R script.

To find all DNC transcripts in the yeast genome, we first updated the current SacCer3 gene annotation (downloaded from www.yeastgenome.org and supplemented with CUTs and SUTs from (Xu *et al*., 2009) using the TranscriptomeReconstructoR package (https://github.com/Maxim-Ivanov/TranscriptomeReconstructoR) and the following previously published datasets: i) Direct RNA-seq from (Garalde *et al*., 2018); ii) CAGE-seq from (Lu & Lin, 2019); iii) 3’ READS from (Liu *et al*., 2017). The borders of known genes were adjusted by the experimental evidence for TSS and PAS called from the CAGE-seq and 3’ READS data, respectively. In addition, 3736 novel transcripts were called from the TSS, PAS, Direct RNA-seq and NET-seq reads which did not overlap with known genes on the same strand. This analysis can be reproduced using the 03-Correct_and_expand_SacCer3_annotation.R script. In effect, we produced a novel gene annotation for *S. cerevisiae* which can be downloaded as a BED file from the code repository.

This improved annotation was used to detect DNC loci, i.e. pairs of nuclear protein-coding genes and non-coding transcripts in divergent orientation with TSSs found within the same nucleosome-free region (NFR) and separated by not more than 500 bp. The search for DNC loci was done using the 04-Find_DNC_loci.R script using MNase data from (Chereji *et al*., 2018) and (Jiang & Pugh, 2009).

### Data deposition

Flow cytometry data of the high-throughput reverse genetic screens were deposited at, FlowRepository, with accession numbers FR-FCM-Z3W4 for the *GCG1*pr screen, and FR-FCM-Z3W5 for the *ORC2*pr screen. NET-seq data were deposited at GEO with the number GSE167499.

### Single-molecule Fluorescence *in situ* Hybridization (smFISH)

21 ml OD_600_ 0.5 cultures were fixed with 4 ml of 32% paraformaldehyde and incubated at RT for 20 min. The fixed cells were washed thrice with 10 ml of ice-cold buffer B (1.2 M sorbitol, 100 mM potassium phosphate buffer, pH 7.5). The fixed cell pellet was resuspended in 0.5 ml of spheroplasting buffer (1.4x buffer B, 0.2% β-mercaptoethanol, 200 mM vanadyl ribonucleoside complex, 300 U lyticase) and incubated at 30°C for 5 min. The resultant spheroplasts were washed and resuspended in 1 ml of ice-cold buffer B. 400 µl of the resuspended cells were spotted on poly-L-lysine coated coverslips and incubated at 4°C, 30 min for the cells to adhere onto coverslips. Later, the coverslips were washed with 2 ml of ice-cold buffer B and incubated in 70% ethanol at −20°C, overnight. For hybridization, the coverslips were placed cell-adhered side on 50 µl drop of hybridization solution (10% dextran sulfate, 10% formamide, and 2x saline-sodium citrate) containing probes labeled with Cy3 and Cy5 (complementary to 14x PP7 and 12x MS2 sequence respectively, 2.5 µM). The coverslips were incubated in a sealed Parafilm chamber at 37°C, 4 hours. After hybridization, the coverslips were washed once with pre-warmed wash buffer (10% formamide, 2x SSC) for 30 min at 37°C, once with 2x SSC, and once with Phosphate buffer saline for 5 min at RT. The rinsed coverslips were air-dried and placed cell-coated side over 15 µl of mounting media with DAPI (ProLong Gold, Life Technologies) on a clean microscope glass slide, allowed to polymerize for at least 24 hours in dark, RT.

#### Imaging

The cells were imaged using Zeiss Axio Observer 7 inverted wide-field fluorescence microscope with LED illumination (SpectraX, Lumencor) and sCMOS ORCA Flash 4.0 V3 (Hamamatsu). A 40x oil objective lens (NA 1.4) with 1.6x Opto var was used. 13 z-stacks were imaged from −3 to 3 µM with 0.5 µM steps and 1×1 binning. An exposure time of 250 ms was used to image Cy3, Cy5, and 25 ms for DAPI channels at 100%, 100%, and 20% LED power, respectively.

#### smFISH data analysis

Image quantification was carried out using a custom Python pipeline. The scripts are available upon request to Lenstra Lab. Images were compressed to 2D images displaying the maximum intensity projection for each pixel across z stacks −3 to 3 µM. Cell and nuclear masks were determined using a custom Python algorithm. Spots corresponding to *GCG1* or *SUT098* transcripts were then counted for cells and nuclei. The transcription site (TS) was defined as the brightest spot in the nucleus and normalized to the median fluorescent intensity of cytoplasmic transcripts. For each sample, three replicate experiments were performed, and approx. 1000 - 3000 cells were counted per strain.

### Single-molecule live-cell imaging

The protocol was followed as described in (Brouwer *et al*., 2020). Yeast cells grown to 0.02 OD_600_ were imaged. The live cells suspended in 4 µl of 2% SC-glucose were spotted on a coverslip and immobilized by placing 2% agarose pads.

#### Imaging conditions

The live cells were imaged using a custom-built microscope consisting of Zeiss Axio Observer 7 inverted wide-field microscope, sCMOS ORCA Flash 4.0 V3 (Hamamatsu), incubator (Okolabs) at 30°C, LED illumination (SpectraX, Lumencor). A 100x oil objective lens (NA 1.46) with 1x optovar and ND filter 1 was used. The images were recorded at 10 s intervals for MS2-*GCG1* and PP7-*SUT098* with 60-time points, 9 z-stacks (−2 to 2 µm) with 0.5 µm steps, 2×2 binning and 200 ms exposure, 30% LED power using micro-manager software.

#### Live-cell image analysis

Maximum intensity projections of the recorded images were computed using Image J Fiji software. A custom-made Python algorithm from Lenstra Lab was used to determine the intensity of the transcription site (TS) for each channel. The algorithm fits a 2D Gaussian mask after local background subtraction and marks the cell boundaries. It tracks the TS overtime in the recorded movies and the output data was manually checked for proper localization of tracking. To determine the on and off periods, a threshold was applied to background subtracted traces of 8 times the standard deviations of the background (for SUT098 transcript). This number was chosen to reliably distinguish on and off periods from background levels at the single transcript level. The scripts are available upon request to Lenstra Lab.

## ACKNOWLEDGEMENTS

We thank the members of Marquardt Lab and Lenstra Lab for discussions and technical assistance. The Marquardt Lab acknowledges the funding from the Novo Nordisk Foundation NNF15OC0014202 and Copenhagen Plant Science Centre Young Investigator Starting grant. This project received support from the European Research Council (ERC) under the European Union’s Horizon 2020 Research and Innovation Programme StG2017-757411 (S.M) and StG2017-755695 (T.L.L), the Netherlands Organization for Scientific Research (NWO, gravitation program CancerGenomiCs.nl, T.L.L.), and Oncode Institute (T.L.L.), which is partly financed by the Dutch Cancer Society. U.G was supported by the European Molecular Biology Organization Short-Term Fellowship (STF-8335) to perform single-molecule studies at the Lenstra Lab.

## AUTHOR CONTRIBUTIONS

S.M and U.G conceived the project with input from all the authors.

U.G, I.S, and N.A.M performed the experiments

M.I, U.G, and D.G.P optimized NET-seq protocol

U.G, N.A.M., I.S, and D.G.P engineered yeast strains

U.G and H.P.P performed the single-molecule imaging

M.I performed the computational analysis of sequencing data and reporter screen.

U.G, T.L.L, H.P.P analyzed and interpreted the single-molecule data

U.G and S.M wrote the manuscript with input from all the authors

## SUPPLEMENTARY TABLES

Table S1. Directionality scores of deletion mutants from *GCG1*pr and *ORC2*pr reporter screen

Table S2. Shortlisted Repressor and Activator mutant candidates from *GCG1*pr and *ORC2*pr reporter screen

Table S3. NET-seq FPKM values of identified transcripts and novel set of DNC loci n=1551

Table S4. List of yeast strains used in the study

Table S5. List of Oligonucleotides used in the study

Table S6. List of Plasmids used in the study

